# AI-Driven Prediction of Drug–Target Interactions for New Chemical Entities

**DOI:** 10.64898/2026.11.18.767501

**Authors:** Vinay Rontala, Bhupender Singh

## Abstract

**Background:** Models that Ranking compounds may improve efficiency in drug discovery; however, due to similar chemistry in train/test sets, performance is often measured using random data splits. Here, we assess the predictive performance on completely novel compounds.

**Method:** Exact Ki values from BindingDB v202609 were used to create a dataset of 421,968 unique ligand-protein interactions. Interactions were defined as active if Ki ≤ 100 nM (53% active). Fingerprints (MACCS, ECFP4/ECFP6), six physicochemical properties, nine ADMET-AI predicted properties and amino acid composition of proteins were used as features, and interactions were classified with XGBoost. Thresholds were optimised on the validation set. External validation was performed using 1,524 target-compound pairs with no exact match in BindingDB. Compounds were defined as active if pChEMBL ≥ 6.

**Results:** Using all three fingerprints, the best performance was achieved with ECFP6 (ROC-AUC 0.9125 [95%CI: 0.9105-0.9145], PR-AUC 0.9222, MCC 0.6613) on 84,394 hidden interactions. Performance dropped significantly on external validation with novel compounds. The lowest drop was seen with ECFP6, with ROC-AUC 0.8040 and MCC 0.4701. All three fingerprints showed a decrease in performance. Performance decreased significantly (by ∼0.11 ROC-AUC and ∼0.19 MCC) when applied to novel compounds compared to performance in random splits. Uncertainty in performance is much less (∼±0.02).

**Conclusion:** Gradient boosting using fingerprints retains some predictive performance in novel compounds. However, performance in random splits overestimates performance in external validation. This is due to several factors including chemical novelty, mismatch between classes and mismatch between ADMET features in train/validation sets. This is discussed

## 1. Introduction

Drug discovery is very slow and inefficient, with only a small amount of candidate compounds progressing to clinical trials (Hay et al., 2014). It can take between 1 and 8 years to discover drugs (U.S. Food and Drug Administration, n.d.), and so methods to prioritise compounds for further experimentation may offer improvements (Paul et al., 2021, Vamathevan et al., 2019).

DTI prediction methods have been developed to predict which compounds are likely to bind to what proteins based on known binding affinities. These methods vary from similarity based gradient boosting approaches (Bleakley & Yamanishi, 2009; He et al., 2017; Pahikkala et al., 2015) to deep learning approaches (Bai et al., 2023; Huang et al., 2021; Nguyen et al., 2021; Öztürk et al., 2018). Even when performing well, it is difficult to interpret model performance. The cutoff for activity may bias results and is often neglected. Random data splitting leads to highly similar analogues in train/val/test sets, over estimating performance in novel chemical types (Sieg et al., 2019; Wu et al., 2018). Additionally, accuracy may give misleading indications in imbalanced datasets. PR-AUC and MCC are preferable metrics (Chicco & Jurman, 2020; Saito & Rehmsmeier, 2015).

Here, a transparent Ki based prediction model was created. (i) Activity cut off was considered, (ii) Models were compared using the same framework for different fingerprints, (iii) Physicochemical properties and nine predicted ADMET properties were included as features, and (iv) models were validated on novel chemical entities with no exact match in BindingDB. Uncertainty in model performance is described, and differences supported and not supported by the data.

## 2. Materials and Methods

### 2.1 Data and labels

Data was extracted from BindingDB_All_202609, in batches of 100,000 rows. Ligand SMILES, target names, protein sequences from first chain, and Ki (nM) were recorded. Any rows with missing data were removed, along with any values with qualifiers (“>10000” etc). 541,854 interactions remained. SMILES were canonicalised using RDKit (Landrum, n.d.), and any molecules that could not be parsed were removed. Duplicate interactions were removed according to canonical SMILES and protein sequences, leaving 421,968 interactions. Interactions were considered active if Ki ≤ 100 nM. Using a prototype of 406,789 interactions, stricter activity cut-offs were investigated. As fewer actives were produced (Table 1), an activity cut-off of 100 nM was selected to create a balanced problem. This resulted in 54.6% active interactions in the cleaned dataset and 53.1% in the final test set, to stabilise F1 and MCC

**Table 1.** Proportion of actives at different Ki cut offs.

| Ki cut-off | Active | Inactive | Proportion of Actives |
|---|---|---|---|
| < 10 nM | 111,282 | 295,507 | 27.4% |
| < 30 nM | 156,806 | 249,983 | 38.5% |
| < 50 nM | 180,142 | 226,647 | 44.3% |
| **≤ 100 nM (adopted; 541,854 rows)** | **295,855** | **245,999** | **54.6%** |

### 2.2 Features

Four sets of features were included for each interaction. For ligands, MACCS (167 bits) (Durant et al., 2002), or ECFP4/ECFP6 (2,048 bits) fingerprints were included. The Morgan radius was 2/3 for the two ECFPs respectively (Rogers & Hahn, 2010). Physicochemical features were included: molecular weight, Crippen logP, TPSA, hydrogen bond donors, hydrogen bond acceptors and QED (Bickerton et al., 2012). Nine ADMET-AI features (Swanson et al., 2024) were included (Caco-2 permeability, intestinal absorption, plasma protein binding, CYP3A4/CYP2D6 inhibition, hepatocytes clearance, half life, hERG and DILI). Missing values were included with a flag, and the median of training data imputed. Lastly, 20 dimensional amino acid composition (AAC) of proteins were included. ADMET-AI was run in batches on 233,386 unique SMILES. Features were predicted values rather than measurements. Resulting matrices contained 211 features for MACCS and 2,092 for ECFP fingerprints.

### 2.3 Model creation, splitting and evaluation

XGBoost (Chen & Guestrin 2016) with 1,500 trees, learning rate 0.03, depth 8, subsampling rate 0.8 and class weights were used. A stratified random split (seed 42) was made to create 20% test data (84,394 interactions total, 44,824 active, 39,570 inactive). 10% of remaining interactions were used to create early stopping points and select thresholds (approx. 303,800 training interactions, approx. 33,800 validation interactions). Since pairs were used, the same ligand/protein may be present in both train and test. Thresholds ranging from 0.30-0.68 were tested on validation data. Thresholds were selected as those with the highest validation accuracy. Thresholds were 0.48 for MACCS and ECFP4, and 0.46 for ECFP6. Atom Pair (Carhart et al., 1985) and Topological Torsion (Nilakantan et al., 1987) fingerprints were also tested under the same framework, for comparison to MACCS. To compare to baselines, a random forest model using MACCS+AAC was run alongside XGBoost using ECFP4+AAC, using a strict activity cut-off (Ki < 10 nM).

Performance is given in ROC-AUC, PR-AUC, accuracy, F1 and MCC. Uncertainty in performance within internal validation was calculated. 95% intervals were calculated using the Hanley-McNeil method in ROC-AUC (Hanley & McNeil, 1982), and using a multinomial bootstrap method (20,000 samples) in threshold metrics. Only uncertainty in sampling from the test data is given, therefore comparisons between fingerprints are purely descriptive. Analyses were carried out using Python, with RDKit, XGBoost, scikit-learn and ADMET-AI.

### 2.4 External validation with NCEs

A table of NCEs with ChEMBL IDs and pChEMBL values was curated (Mendez et al., 2019). 7,319 NCEs were valid. 1,057 interactions with canonical SMILES present in BindingDB were removed, leaving 6,262 NCEs. If compounds had pChEMBL values, they were mapped to a target and UniProt sequence (UniProt Consortium, 2023) (99 targets in total). If pChEMBL ≥ 6 the compound was considered active (≤ 1 µM). From 1,730 unique NCEs, all compound-target interactions (1,524) with these data were used for external validation. There was no ADMET data available for these interactions, so ADMET values were imputed with medians from training, and all missing value flags were set to 1. Models were scored using validation thresholds. As exact matching was used, similar analogues may still be present in external validation. Additionally, whether the targets in external validation overlapped with training data was not assessed. Therefore, this is not scaffold or target held out validation. Uncertainty in external validation is +/- ∼0.022 for accuracy, and +/- 0.021-0.024 for ROC-AUC (active proportion 0.4-0.7).

## 3. Results

### 3.1 Internal performance

All models performed well. For 84,394 hidden interactions, ECFP6 performed best with all metrics (Table 2, Figure 2). Differences between model performances were small (differences in ROC-AUC 0.0050, PR-AUC 0.0064, MCC 0.0090). ECFP6 predicted the most true positives (38,480) and the least false negatives (6,344), giving high sensitivity and specificity (sensitivity 0.859, specificity 0.801). MACCS and ECFP4 performed with slightly lower sensitivity (0.849, 0.841) and specificity (0.802, 0.813). AUC increased up to 1,500 trees (0.9056, 0.9075, 0.9115), meaning models had not fully converged. Atom Pair and Topological Torsion performed worse than MACCS and were not taken forward (see Supplementary Table S1). In the baseline comparison, XGBoost performed better than random forest in ROC-AUC (0.9195 vs. 0.9084) and recall (0.8309 vs. 0.7164). However, precision decreased (0.6836 vs. 0.7440). Therefore, XGBoost was selected.

**Table 2.** Performance in internal validation of BindingDB. 95% confidence interval in brackets.

| **Fingerprint** | **Threshold** | **ROC-AUC** | **PR-AUC** | **Accuracy** | **F1** | **MCC** |
|---|---|---|---|---|---|---|
| MACCS | 0.48 | 0.9075<br>[0.9055-0.9095] | 0.9158 | 0.8270<br>[0.8245-0.8296] | 0.8391<br>[0.8365-0.8417] | 0.6524<br>[0.6472-0.6576] |
| ECFP4 | 0.48 | 0.9093<br>[0.9073-0.9113] | 0.9189 | 0.8280<br>[0.8254-0.8305] | 0.8386<br>[0.8360-0.8412] | 0.6545<br>[0.6494-0.6596] |
| **ECFP6** | **0.46** | **0.9125**<br>**[0.9105-0.9145]** | **0.9222** | **0.8315**<br>**[0.8289-0.8340]** | **0.8440**<br>**[0.8415-0.8465]** | **0.6613**<br>**[0.6562-0.6664]** |

**Figure. 1.**
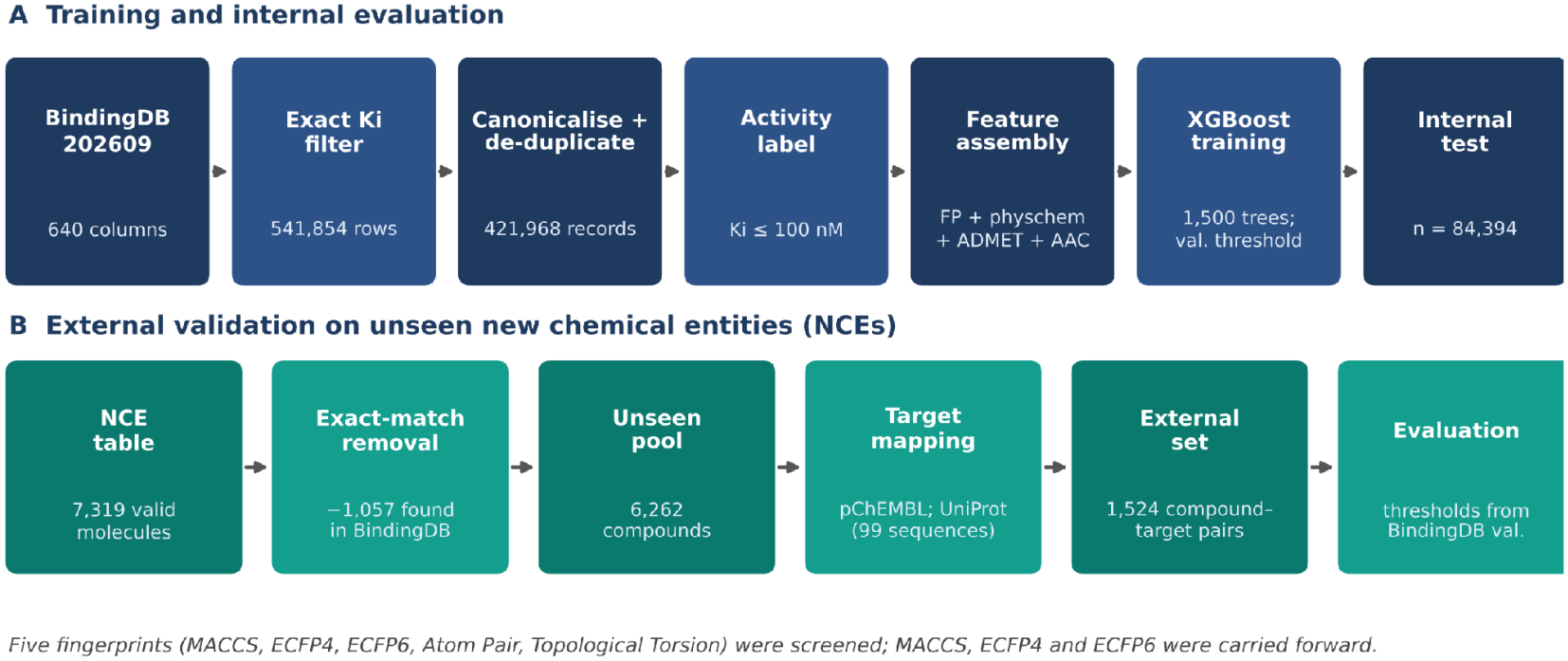
Study design: (A) Curation, features, training and internal validation; (B) construction of the external NCE s.

**Fig. 2.**
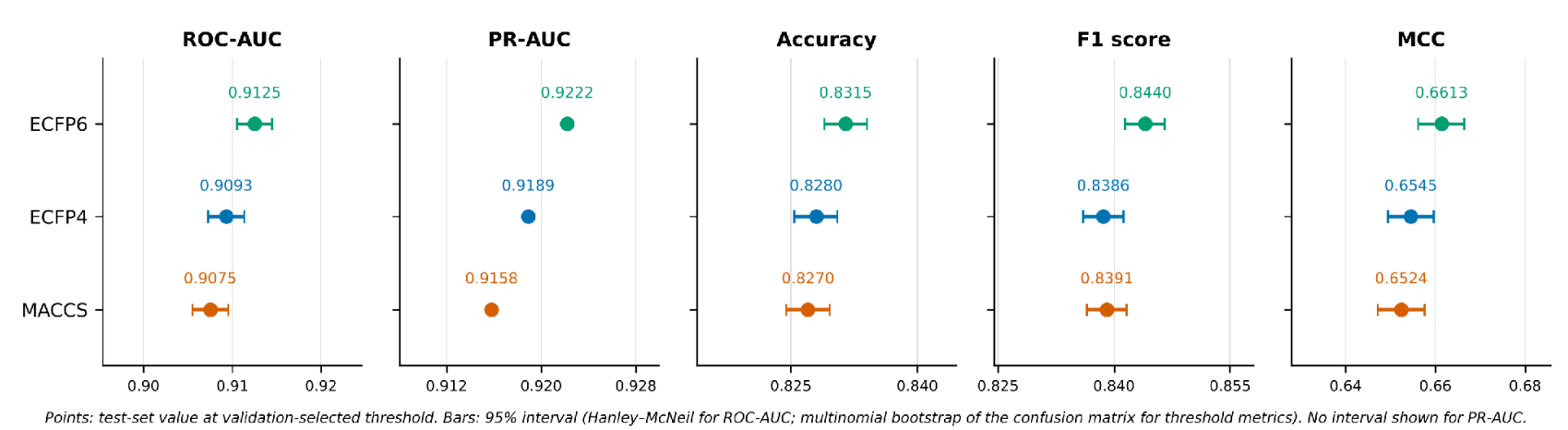
Internal validation performance with 95% confidence intervals.

### 3.2 External validation & performance drop

Performance dropped significantly in external validation. ECFP6 gave the best performance (Table 3, Figure 3), with ROC-AUC of 0.8040 and MCC of 0.4701. ECFP6 had the smallest drop in all metrics compared to MACCS and ECFP4. Within external validation, ROC-AUC dropped by 0.1085 (ECFP6) compared to 0.1180 (MACCS) and 0.1209 (ECFP4). MCC dropped by 0.1912 (ECFP6) compared to 0.2227 (MACCS) and 0.2381 (ECFP4). Compared to uncertainty in sampling during internal validation, there is a large drop in performance in external validation. However, differences between models in external validation are small (∼0.016 difference in ROC-AUC). Therefore, the ordering of models in external validation is suggestive only. In a pilot analysis with 228 pairs (ECFP4), ROC-AUC was 0.735 and recall of actives was 0.47. However, the large number of pairs (1,524) gives a more reliable measure.

**Table 3.** Performance on 1,524 novel NCE-target interactions compared to performance in BindingDB testing. Differences in performance are shown in brackets.

| **Fingerprint** | **Threshold** | **ROC-AUC** | **PR-AUC** | **Accuracy** | **F1** | **MCC** |
|---|---|---|---|---|---|---|
| MACCS | 0.48 | 0.7895<br>[−0.1180] | 0.8232<br>[−0.0926] | 0.7106<br>[−0.1164] | 0.7249<br>[−0.1142] | 0.4297<br>[−0.2227] |
| ECFP4 | 0.48 | 0.7884<br>[−0.1209] | 0.8302<br>[−0.0887] | 0.7041<br>[−0.1239] | 0.7187<br>[−0.1199] | 0.4164<br>[−0.2381] |
| **ECFP6** | **0.46** | **0.8040**<br>**[−0.1085]** | **0.8464**<br>**[−0.0758]** | **0.7316**<br>**[−0.0999]** | **0.7464**<br>**[−0.0976]** | **0.4701**<br>**[−0.1912]** |

**Figure 3.**
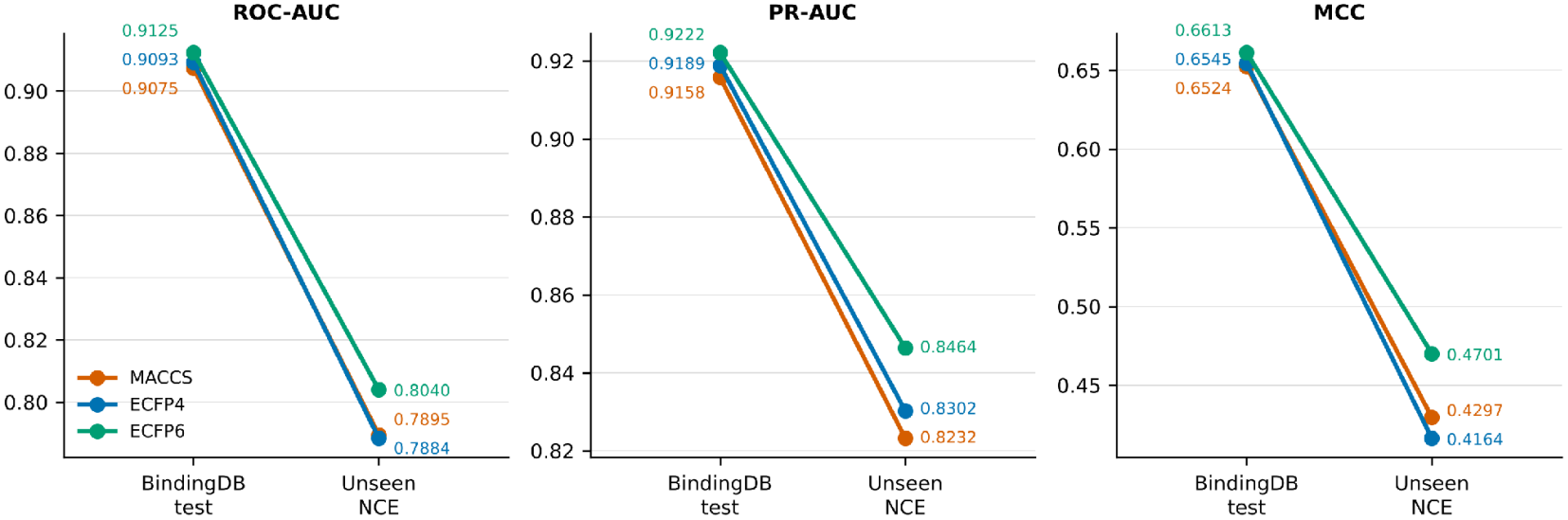
Model performance in internal validation (BindingDB) and external validation (novel NCEs).

## 4. Discussion

Within BindingDB, all models have good performance, with ECFP6 having the highest ROC-AUC (0.91) and MCC (0.65-0.66). When comparing MACCS, ECFP4 and ECP6 performance in a random split, there was very little difference between models. The 167 bit MACCS keys performed very similarly to the 2048 bit circular fingerprints (difference < 0.005 ROC-AUC) and larger Atom Pair and Topological Torsion vectors did not perform better. MACCS therefore provides a valid efficient option. However, performance decreased significantly in external validation with novel NCEs. ECFP6 decreased by ∼0.11 ROC-AUC and 0.19 MCC.

This performance drop in external validation is due to a combination of factors. (1) Chemical novelty – using a random data split, structurally similar analogues, ligands and proteins may appear in both the train and test sets, leading to increased performance in internal validation. (2) Class shift – during training, Ki were ≤ 100 nM, whilst external data had pChEMBL ≥ 6 (≤ 1 µM) across a collection of assays. This would reduce performance in accuracy, F1 and MCC much more than ROC-AUC, as seen. A re-evaluation with matched thresholds would provide a direct indication of this. (3) ADMET features – in external validation, ADMET features were imputed and all flags set, which was rarely seen in training data. Therefore, it is unknown whether specific ADMET information for compounds helps in predicting interactions with novel compounds. (4) Coarse features – AAC cannot differentiate between similar protein targets. ECFP6 performed best in external validation, suggesting that larger atomic environments have better generalisability to novel scaffolds. However, this is a trend and not prove

In practice the model should reorder experimental work: score new analogues against a target, assay those above the validated threshold with acceptable predicted ADMET, and deprioritise the rest. An external ROC-AUC of 0.804 means an active compound outranks an inactive one about 80% of the time. Predictions are most defensible within the training domain (Sahigara et al., 2012; Tropsha, 2010); we implemented no applicability-domain score and measured no time or cost savings. Limitations also include a single seed and split, no feature ablations, and external class prevalence that must be reported with PR-AUC. Next steps are matched-label re-evaluation, ADMET-AI predictions for the NCEs, scaffold- and target-held-out splits, ablations, multiple seeds, learned protein embeddings (Lin et al., 2023), calibration, and prospective assay testing.

## 5. Conclusions

Reproducible methods were used to create a dataset of 421,968 unique Ki values from BindingDB. Different fingerprints were compared using the same framework, alongside physicochemical and predicted ADMET features. ECFP6 had best performance in internal validation (ROC-AUC 0.9125, MCC 0.6613) on 84,394 hidden interactions. In external validation using 1,524 novel NCE-target interactions, ECFP6 had a ROC-AUC of 0.8040, PR-AUC of 0.8464 and MCC of 0.4701. Therefore,Whilst ranking performance is retained on novel chemical entities, performance decreases by ∼ 0.11 ROC-AUC and ∼0.19 MCC compared to performance in random data splits. Performance in external validation, with uncertainty, gives the realistic expectation for prospecting.

## Supporting information

https://docs.google.com/document/d/1BsdGcIpz1G5E4Icg74zDre7El-eEhySbfwwXrY9ysXo/edit?usp=sharing

## Supplementary Table S1

**Table S1.** Performance in testing from BindingDB. All fingerprints were tested under the same framework.

| **Fingerprint** | **Bits** | **ROC-AUC** | **PR-AUC** | **Accuracy** | **F1** | **MCC** |
|---|---|---|---|---|---|---|
| MACCS | 167 | 0.9075 | 0.9158 | 0.8270 | 0.8391 | 0.6524 |
| ECFP4 | 2048 | 0.9093 | 0.9189 | 0.8280 | 0.8386 | 0.6545 |
| ECFP6 | 2048 | 0.9125 | 0.9222 | 0.8315 | 0.8440 | 0.6613 |
| Atom Pair | 2048 | < 0.9075 | < 0.9158 | < 0.8270 | < 0.8391 | < 0.6524 |
| Topological Torsion | 2048 | < 0.9075 | < 0.9158 | < 0.8270 | < 0.8391 | < 0.6524 |

*Atom Pair and Topological Torsion performed worse than MACCS in all metrics. Values provided for MACCS are therefore upper limits..*

---

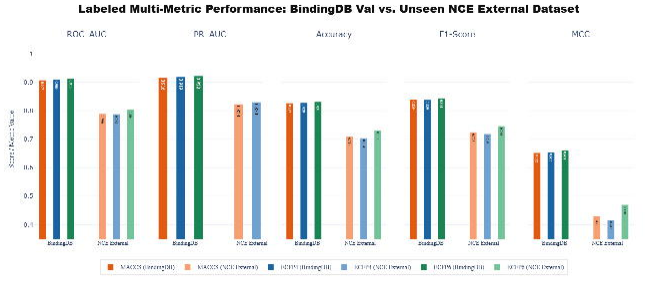

