## Supplementary material for "AI-Driven Prediction of Drug–Target Interactions for New Chemical Entities": https://docs.google.com/document/d/1BsdGcIpz1G5E4Icg74zDre7El-eEhySbfwwXrY9ysXo/edit?usp=sharing

#### A Project Report on

**Submitted By:**
Student Name : Vinay Rontala
Course Name : AIML in DD
Project Duration : 6 Months

**PI Details:**
Dr Bhupender Singh
CRO Scientist
Biotecnika Info Labs Pvt Ltd

**November 2026**
(Month, Year of Submission )

#### Declaration by the candidate

I, '**Vinay Rontala**', hereby declare that the project work entitled **"AI-Driven Prediction of Drug-Target Interactions for New Chemical Entities"** submitted to Biotecnika Info Labs is an original work carried out by me under the guidance of Dr Bhupender Singh.

I declare that this project has not been submitted to any other University or Institution for the award of any degree or diploma or any other title.

Further, I declare that I shall not publish, present, or communicate any part of this project work in any form — including journals, conferences, or digital media — without the prior written consent of my Principal Investigator and the host organization.

I understand that failure to obtain such prior approval shall be treated as a breach of institutional and research ethics, and may result in disciplinary actions as deemed appropriate by the organization.

By signing below, I acknowledge that I have read and agree to abide by the above statement.

Student Signature :
Student Name : Vinay Rontala
Course Name : AIML in DD
Project Duration : 6 Months

Dr Bhupender Singh
Scientist & Trainer
Biotecnika Info Labs Pvt Ltd

### Acknowledgement

I express my sincere gratitude to my project investigator, Prof. Dr. Bhupeder Singh, for his continuous guidance, encouragement and constructive criticism throughout this project. His insight into drug discovery and computational pharmacology shaped both the scientific questions I asked and the care with which I tested our answers.

I am also grateful to the whole Biotecnika team for providing the learning environment, resources and support that made this work possible. The discussions, feedback and the opportunity to apply bioinformatics and machine learning to a real drug discovery problem have been invaluable.

Finally, I thank for their patience and support.

**Vinay Rontala**

### Contents

**Acknowledgement** .................................................. **2**

**Contents** .................................................. **3-4**

**List of Figures** .................................................. **5-6**

**List of Tables** .................................................. **7-8**

**Abstract** .................................................. **9**

**1. Introduction**.................................................. **10-16**

1.1 The Drug Development Lifecycle.................................................. 11-12
1.2 Clinical Trial Phases.................................................. 12-13
6.4 Future Scope.................................................................................................................... 42-43
**References....................................................................................................... 44-46**

### List of Figures

| FIGURE NO. | DESCRIPTION | PAGE NO |
|---|---|---|
| **Figure 1** | Attrition funnel from 10,000 candidates to one approved drug (left) and typical duration of each development stage (right) | **1** |
| **Figure 2** | End-to-end workflow of the project with the record counts at each stage | **7** |
| **Figure 3** | Share of active compounds under each Ki cut-off | **8** |
| **Figure 4** | Comparison of MACCS, ECFP4 and ECFP6 across the five evaluation metrics on the BindingDB test set | **11** |
| **Figure 5** | Performance gaps between the three fingerprint models, with exact scores and the difference zone | **11** |
| **Figure 6** | Confusion matrices of the MACCS, ECFP4 and ECFP6 models at their validation-selected thresholds. | **12** |
| **Figure 7** | Point-plot comparison of BindingDB validation and unseen NCE performance for each fingerprint | **13** |

### List of Figures

| | | |
|---|---|---|
| **Figure 8** | Labelled multi-metric performance: BindingDB test versus unseen NCE external dataset | **13** |

### List of Tables

| TABLE NO. | DESCRIPTION | PAGE NO |
|---|---|---|
| **Table 1** | Drug development stages and typical durations | **1** |
| **Table 2** | AI models mapped to drug discovery tasks | **3** |
| **Table 3** | Fields retained from the BindingDB master file | **7** |
| **Table 4** | Effect of the Ki cut-off on class balance | **8** |
| **Table 5** | Feature blocks used for model training | **9** |
| **Table 6** | Random forest versus XGBoost baselines | **9** |
| **Table 7** | Performance on the BindingDB held-out test set | **11** |
| **Table 8** | Confusion-matrix results on the BindingDB test set | **12** |

### List of Tables

| TABLE NO. | DESCRIPTION | PAGE NO |
|---|---|---|
| **Table 9** | Model comparison of the tested fingerprints, ranked by performance | **13** |
| **Table 10** | Performance on the unseen NCE external set (1,524 pairs) | **12** |
| **Table 11** | Drop in performance from the BindingDB test set to the NCE set | **14** |
| **Table 12** | Proposed next steps | **16** |

### Abstract

We built an end-to-end machine-learning pipeline that predicts whether a small molecule binds a protein target potently, using inhibition constants (Ki) from BindingDB. The raw BindingDB release (202609) was filtered to 541,854 exact Ki measurements, canonicalised with RDKit and de-duplicated to 421,968 unique ligand–protein records. Ki values were binarised at 100 nM (active ≤ 100 nM), a cut-off that gives a nearly balanced dataset. Each record was encoded with a ligand fingerprint, six physicochemical descriptors, nine ADMET-AI properties and the 20-dimensional amino-acid composition of the target, and was classified with XGBoost.

Five fingerprints (MACCS, ECFP4, ECFP6, Atom Pair and Topological Torsion) were compared; Atom Pair and Topological Torsion scored below MACCS, so MACCS, ECFP4 and ECFP6 were studied in full. On an 84,394-record held-out BindingDB test set, ECFP6 performed best (ROC-AUC 0.9125, PR-AUC 0.9222, accuracy 83.15%, F1 0.8440, MCC 0.6613), although all three fingerprints were within 0.005 ROC-AUC of each other. On an external set of 1,524 NCE–target pairs absent from BindingDB, ROC-AUC fell to 0.788–0.804 and MCC to 0.42–0.47, with ECFP6 degrading least on every metric. We analyse the causes of this generalisation gap, describe how the model supports early drug discovery and outline future work.

**Keywords:** drug–target interaction, BindingDB, molecular fingerprints, XGBoost, ADMET-AI, external validation.

### 1. Introduction

### 1. Introduction

Drug discovery is a highly complex and lengthy process which aims to find small molecule drugs which can interact with relevant proteins involved in causing a particular disease. This book will provide insight into the various stages of drug development giving an overview of the key steps involved in drug discovery prior to clinical trial. The molecular and biological concepts behind drug-target interactions will be covered, alongside showing how computational techniques and AI are playing a greater role in drug discovery. Finally the motivation for undertaking this project will be discussed.

#### 1.1 The Drug Development Lifecycle

The pharmaceutical pipeline is a multi-stage funnel designed to isolate one viable therapeutic from a very large library of candidates. Historically, the process begins with approximately 10,000 initial compounds. Only about 250 progress to pre-clinical testing, a mere 5 enter clinical trials, and ultimately a single compound achieves the safety and efficacy profile required for approval by the U.S. Food and Drug Administration (FDA) (Figure 1).

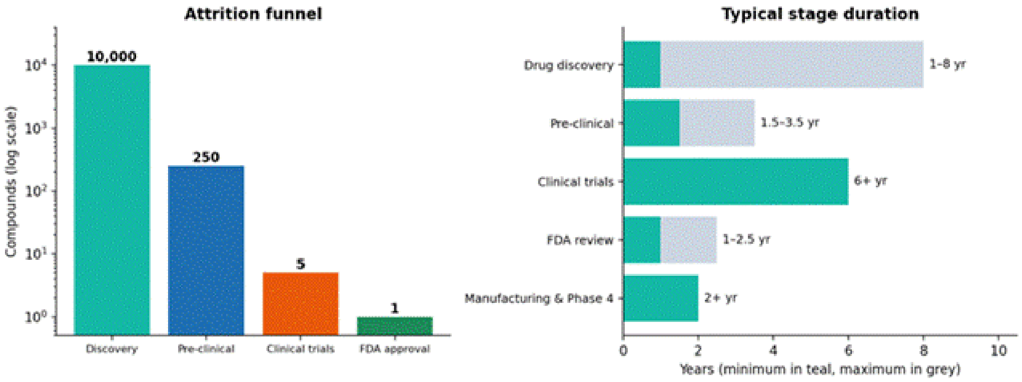

**Figure 1.** Attrition funnel from 10,000 candidates to one approved drug (left) and typical duration of each development stage (right).

Moving from a computational hit to a marketed drug is a decade-scale endeavour. Table 1 presents the approximate time involved in each stage of drug development, based on industry benchmarks. The discovery phase itself may take as long as eight years. During this period, a large number of chemical compounds are examined, but only a small proportion are considered suitable for further development. Most candidates are eliminated because they do not show the required activity or other desirable properties. This makes early-stage computational screening particularly useful, as it can help prioritise promising compounds before investing time and resources in laboratory experiments.

**Table 1.** Drug development stages and typical durations

| Stage | Typical Timeframe | Key activities |
|---|---|---|
| Drug discovery | 1 – 8 years | Target identification/ hit discovery/ lead optimisation |
| Pre-clinical testing | 1.5 – 3.5 years | In vitro & Animal Models; pharmacology, safety & formulation development |
| Clinical Trials | 6+ years | Human testing to assess safety profile, dosing and confirm efficacy |
| FDA review | 1 – 2.5 years | Analysis of the New Drug Application (NDA) , safety data |
| Manufacturing and Phase 4 | 2+ years | Large-scale manufacturing and post-market surveillance |

The discovery stage alone can take up to eight years, and it is also where the largest number of compounds is screened and discarded. This is why any method that ranks candidates reliably before laboratory work begins has high value.

#### 1.2 Clinical Trial Phases

Once a candidate has completed preclinical evaluation and an Investigational New Drug (IND) application has been submitted, it may proceed to clinical testing in humans. Clinical development is generally divided into three main phases:

- **Phase I:** This phase of clinical testing will look mainly at the safety, tolerability and pharmacology, typically using either 20-80 healthy volunteers.
- **Phase II:** The focus shifts towards evaluating the drug's effectiveness, dose response and short-term safety. Studies generally include approximately 100–200 participants.

- **Phase III:** Larger clinical studies are conducted to confirm therapeutic efficacy, compare outcomes and monitor adverse reactions. These trials may involve 1,000–10,000 or more participants.

These stages help find out if there is enough evidence for a candidate to be progressed further for regulatory assessment.

### 1.3 The Drug Discovery Workflow:

#### From Hit Identification to Lead Optimisation

Drug discovery consists of a series of interconnecting processes. First, a suitable biological target is identified.

**Target identification & validation:** A biological element, such as proteins, enzymes and receptors, that have a key role within the disease of interest is determined. Evidence is sought to provide biological proof of involvement in the disease process.

**Hit identification:** Once a target has been identified, compounds are screened for those that can interact with this target. Ligand-based methods can be employed to screen hits. Molecular fingerPrinting, Pharophore Searching and Scaffold Hopping methods take advantage of the similarities of the fingerprints of the compounds. Structure-based methods include Protein Conformation Analysis and Molecular Docking methods.

**Lead optimisation:** Leads with promising activity are modified to increase their potency, selectivity and safety. Typically this occurs using an iterative Design-Make-Test-Analyse cycle (DMTA). Structure Activity Relationship (SAR) studies are performed to understand how changes to the compound's structure can affect its biological activity and inform the selection of improved leads.

This project focuses predominately on hit identification and initial lead profiling. Machine learning methods are applied to predict the potential interaction between drugs and targets. Prediction of ADMET properties increases the knowledge of the compounds by providing information on their pharmacokinetic profile and safety.

#### 1.4 Molecular Representations and Cheminformatics

Before processing molecules for computational analysis they must be translated into a format that can be read by computers. A commonly used format is the Simplified Molecular Input Line Entry System (SMILES). For example, ethanol can be represented as CCO whilst benzene would be c1ccccc1.

This is a concise format for representing chemical molecules and is beneficial for storing information relating to large molecular databases. This may involve processing millions of molecules. This commonly used format is beneficial for processing chemical structures retrieved from PubChem and many other compound databases.

This project involved using RDKit. RDKit is an open source cheminformatics toolkit. Using RDKit it is possible to generate molecular objects from SMILES strings, draw the molecule in 2D and compute key physicochemical properties. Physicochemical properties include molecular weight (MW), lipophilicity (LogP) and topological polar surface area (TPSA). RDKit allows the creation of molecular fingerprints. This is possible using the Morgan fingerprint tool in RDKit which gives Extended-Connectivity Fingerprints (ECFP). The molecular representations provided will give chemical features that can be used for machine learning predictions.

#### 1.5 Protein Targets and Drug-Binding Mechanisms

Proteins play vital roles in the human body, such as cell signalling, metabolism and immune regulation. Many drugs exert their activity by specifically targeting proteins. Some common drug targets include enzymes such as kinases, G-protein coupled receptors (GPCRs), ion channels and transport proteins.

When thinking about molecular binding of compounds to proteins, there are two commonly discussed concepts. First is the lock and key theory where the ligand fits into a complementary binding pocket. The second is the induced-fit theory which takes into consideration the flexible nature of proteins and how they may change conformation to allow binding of the ligand.

The strength of the binding between a drug and a protein can be described using binding affinities. This can depend on several factors, including hydrogen bonding, hydrophobic interactions, ionic interactions and van der Waals forces. These are often expressed using values such inhibition constant (Ki), dissociation constant (Kd) and half maximal inhibitory concentrations (IC50). Overall lower values suggest stronger binding or higher inhibitory potency. However these values represent different experimental values and should not be considered equivalent.

This project uses Ki values obtained from BindingDB as the basis for evaluating compound–target interactions and constructing the classification dataset

#### 1.6 Artificial Intelligence in Modern Pharmacology

The increasing usage of artificial intelligence means that new methodologies have been developed throughout multiple stages of drug discovery. Often traditional drug discovery methodologies involve numerous time-consuming lab based experiments, with the resulting increase in development times and costs. AI techniques provide another method for analysing data from chemical and biological processes to help observe patterns.

Some important applications include drug–target interaction prediction, early toxicity assessment, drug repurposing and the design of new molecular structures. Machine-learning models can examine molecular fingerprints, chemical descriptors and protein sequence information to estimate the likelihood of interactions between compounds and biological targets.

In the present work, AI-based prediction is applied to drug–target interaction analysis, with additional ADMET information used to support early assessment of candidate compounds. Table 2 presents an overview of commonly used AI models and their applications across different drug discovery tasks.

**Table 2.** AI models mapped to drug discovery tasks

| Discovery Process | Design Focus | AI model |
|---|---|---|
| Target identification | Prediction of PPI, protein folding | Analysis of druggability sites (FD/DCA), Prediction of residues (CNN) |
| Hit discovery | Virtual Screening, Scoring Activity, Drug Repurposing | SVM, Adversarial Autoencoders, Network Pathology |
| Hit-to-Lead | QSAR, De Novo Design | Traditional ML, DNN, Deep Reinforcement Learning |
| Lead optimisation | Evaluation of ADMET/ property | CNN, RNN, Multi-Task Neural Networks |

A DTI model takes the ligand (SMILES) and the protein (FASTA sequence) as inputs. The ligand is converted into fingerprints with RDKit and the protein into numerical descriptors. Candidates are also assessed using important physicochemical properties that help determine their drug-like characteristics. These include molecular weight (MW), LogP, hydrogen bond donors and acceptors, topological polar surface area (TPSA), and toxicity-related parameters. Different machine learning approaches are used to analyse these molecular features. Traditional methods, such as Random Forest and Support Vector Machines (SVM), are commonly applied in classification and quantitative structure–activity relationship (QSAR) studies. More advanced approaches, including Convolutional Neural Networks (CNNs), Recurrent Neural Networks (RNNs), and Graph Neural Networks (GNNs), can identify more complex patterns in molecular structures. Depending on the objective, the model may predict whether a drug interacts with a particular target (binary classification) or estimate the strength of that interaction through a binding-affinity score (regression).

#### 1.7 Motivation for This Work

Despite having a large quantity of data available, the public databases often have differences in data quality. The manner in which data is collected, filtered and annotated, as well as how it is separated for training and testing, can have a big impact on results reliability.

This study focuses on creating a transparent and systematic prediction pipeline. Various molecular fingerprint methods were used with the Ki values from BindingDB. They were investigated with the same experimental conditions. Physiochemical descriptor and ADMET properties were added to give insight into the data. The models were created and tested with new chemical entity data, outside of those found in BindingDB.

### 2. Review of Literature

### 2. Review of Literature

#### 2.1. Binding Affinity Datasets

Prediction of drug-target interactions (DTI) relies heavily on binding affinity information obtained from experimental data. BindingDB is a publicly available binding affinity database that gives quantitative values for protein-ligand interactions such as inhibition constant (Ki), dissociation constant (Kd), half maximal inhibitory concentration (IC50) and half maximal effective concentration (EC50) *(Liu et al., 2007; Gilson et al., 2016)*. This dataset is very useful for developing larger machine learning models due to the large amount of bioactivity data. However, this dataset contains experimental data under various conditions.

Choosing a suitable binding affinity value is important as some types are more consistent. Ki and Kd are equilibrium values for inhibitor binding and protein-ligand complex formation. IC50 indicates the concentration of compound needed to give 50% inhibition in an assay. This depends on the experimental setup of the assay as it is dependent on substrate concentration and enzyme kinetics. Assuming suitable conditions, IC50 values can be converted into Ki values using the Cheng-Prusoff equation *(Cheng and Prusoff, 1973).*

Due to the dependence on experimental conditions, Ki values were prioritised in this study.

Other datasets can also be used to characterise molecules. ChEMBL is a manually curated dataset containing bioactivity data obtained from medicinal chemistry data *(Mendez et al., 2019)*. Data can be scaled to a uniform pChEMBL scale using the negative logarithm of potencies in M. This allows multiple different activity values such as Ki, Kd and IC50 to be compared. ChEMBL will be used to characterise the external NCE dataset.

PubChem contains standardised molecular structures, compound information and biological screening data *(Kim et al., 2023)*. DrugBank contains information relating to approved and investigational drugs, linking them to biological targets *(Wishart et al., 2018)*. These resources can be used to identify compounds, compare molecular structures and find new novel compounds.

#### 2.2. Machine Learning and Deep Learning Approaches For Drug-Target Interaction Prediction

Initial methods of using machine learning to predict drug-target interactions utilise similarity based methods. Kernel regression and boosting techniques utilise similarities between molecular structures and proteins to predict interactions *(Pahikkala et al., 2015; He et al., 2017)*. These models were found to give promising results. However, they do depend heavily on how similar training examples are to the unknown drug-target pairs.

Recent advances in deep learning have allowed more complex molecular and protein representations.

**Some notable models include:**
DeepDTA – Utilises 1D-CNNs to extract features from SMILES and amino acid sequences *(Öztürk et al., 2018)*
GraphDTA – Molecular structures are represented as graphs and utilise graph neural networks *(Nguyen et al., 2021)*
MolTrans – Utilises transformer based representations with interaction modules to find relationships between molecular and protein substructures *(Huang et al., 2021)*
DrugBAN – Utilises bilinear attention networks to learn interactions *(Bai et al., 2023)*

While these models have advanced methods, many well known DTI models were initially tested on benchmark datasets with low target diversity and high kinase representation. This means that their performance on a range of diverse compounds and unknown proteins needs to be investigated.

Complex deep learning models can require large amounts of computing power.
An engineered feature based machine learning model was used using gradient boosted decision trees. This provided efficient computing and a systematic way of comparing different molecular representations. A large dataset of interactions from BindingDB was used to develop the model.

#### 2.3. Molecular Fingerprints and Gradient Boosted Trees

Molecular fingerprints are numerical representations of molecular structures. They contain information about whether certain substructures are present in the molecule. Variable sized molecules can be converted into fixed sized vectors.

MACCS Keys – MACCS fingerprints contain 166 keys describing different molecular substructures. They provide a simple and easily interpretable representation of molecular structures. *(Durant et al., 2002)*

Extended Connectivity Fingerprints (ECFP) – These fingerprints utilise hashing to describe the connectivity of atoms in a molecule. ECFP4 and ECFP6 are equivalent to Morgan fingerprints with radius 2 and 3 bonds. These fingerprints are generated from the molecular structure itself rather than using defined keys. This allows for a wider representation of molecules. *(Rogers and Hahn, 2010)*

Atom Pair Fingerprints – Describes pairs of atoms with their separation.
Topological Torsion Fingerprints – Sequences of four atoms bonded to each other.
Using the correct fingerprint is important to ensure the model can recognise chemical similarity.
Gradient boosted trees are a useful machine learning technique to use for high dimensionality molecular data. XGBoost *(Chen and Guestrin, 2016)* utilises an optimised gradient boosting algorithm. This allows for efficient computation, regularisation and flexibility with different feature types.

XGBoost provides a fast way of combining molecular fingerprints with additional physiochemical and biological features. This makes it suitable for large scale DTI predictions. XGBoost was chosen as the main algorithm.

#### 2.4. ADMET Prediction and Molecular Drug-Likeness

Identifying compounds with high binding affinity to a biological target is not enough to ensure success in drug discovery. The candidate molecules must have good ADMET properties to aid their development as therapeutics.

Important ADMET properties include gastrointestinal absorption, blood brain barrier permeability, metabolic stability, clearance, interactions with cytochrome P450 enzymes, cardiotoxicity and drug induced liver injury.

ADMET-AI is a computational tool containing graph based deep learning models to predict multiple ADMET properties from large chemical libraries. Its performance has been tested using benchmark datasets *(Swanson et al., 2024)*.

Molecules can also be assessed for drug-likeness using well known physiochemical descriptors.
Lipinski's Rule of Five – Guidelines for assessing the drug-likeness of oral compounds based on molecular weight, lipophilicity, hydrogen bond donors and acceptors. *(Lipinski et al., 1997)*
Quantitative Estimate of Drug-likeness (QED) – A combined measure of drug-likeness based on various physiochemical properties. *(Bickerton et al., 2012)*

Six physiochemical and drug-likeness descriptors were combined with nine ADMET prediction endpoints.

These can be combined with molecular fingerprints to allow the model to consider the molecular characteristics and ADMET properties.

#### 2.5. Model Evaluation

Choosing the correct evaluation measures for your model is important to ensure reliability. Accuracy can be misleading when there is a large difference between the number of active and inactive interactions. Additional measures need to be considered.

Precision Recall Area Under the Curve (PR-AUC) is a useful measure to consider when using imbalanced datasets as it considers the relation between precision and recall. *(Saito and Rehmsmeier, 2015)*

The Matthews Correlation Coefficient (MCC) is another robust measure as it uses all information from the confusion matrix (True Positives, True Negatives, False Positives and False Negatives). *(Chicco and Jurman, 2020)*

PR-AUC and MCC were prioritised.

How the data is divided can greatly influence model performance. If random train/test splits are used and the model encounters chemically similar compounds in both the training and test set the model may perform unrealistically well. *(Sieg et al., 2019)*

**More rigorous validation methods include:**

Scaffold Split – Divide compounds by scaffold.
Cluster Split – Group chemically similar molecules.
Target Holdout – Test performance on proteins not included in training.
External Validation – Test model on independent compounds.

External validation is important in computational drug discovery to give a realistic view of model performance. *(Tropsha, 2010)*

The external NCE dataset was split using exact compound removal to ensure minimal overlap with the BindingDB training data.

#### 2.6. Potency, Molecular Properties and Ligand Efficiency

The relationship between molecular structure, physiochemical properties and potency is important in computational drug discovery.

In experimental datasets, binding affinity may increase with increasing molecular size and lipophilicity due to extra interactions with the protein binding pocket. However, this does not necessarily mean that the molecule has good binding efficiency.

Ligand efficiency can be used to overcome this problem by accounting for the size of the molecule when measuring binding potency. This can be used to distinguish between compounds with favourable binding characteristics and those where increased potency is due to increased complexity.

Machine learning models using potency labelling may learn about the relationship between molecular weight, lipophilicity and other properties.

When applying models trained on historical BindingDB compounds to newly optimised NCEs there may be differences in physiochemical distributions.

This can bias predictions and lower performance on external validation. Therefore molecular properties must be considered.

#### 2.7. Activity Cliffs and Limitations of Similarity Based Models

The similarity principle is an important concept in computational chemistry. It states that molecules with similar structures will have similar biological activities.

Many molecular fingerprint based models take advantage of this. However,

This is not always the case. Activity cliffs refer to situations where molecules with similar structures have significantly different biological potencies. This can be caused by small structural changes such as changing a halogen atom, functional group or stereochemistry. *(Maggiora, 2006; Stumpfe and Bajorath, 2012)*

This may be due to interactions at the binding pocket, changes in molecular conformation, steric effects or other factors.

Activity cliffs can be an issue with molecular fingerprint based machine learning models as compounds with similar molecular fingerprint may have very different binding affinities.

This can be an issue when screening NCEs from new chemical series as small changes in structure may affect important interactions with the protein.

Therefore molecular fingerprint based DTI models should be considered along with their applicability domain and external validation performance.

#### 2.8. Protein Representations and Feature Engineering

Protein representation is an important aspect of computational drug-target interaction prediction. Different representations can be used to extract more information.

Amino Acid Composition (AAC) is a common method used to represent proteins. It takes a protein and gives a vector of 20 features describing the relative proportion of each amino acid in the protein.

AAC is fast and easy to implement however it does not give any information about the order of amino acids, interactions between residues, domains or binding site specific information.

This can cause problems as proteins with similar amino acid compositions may have similar representations. This can be an issue when working with closely related protein families such as kinases due to conserved regions and ATP binding sites.

More advanced methods use language models to create contextual representations. The ESM-2 model learns from large protein sequences to identify evolutionary relationships between amino acids. *(Lin et al., 2023)*

This can provide opportunities to gain extra information and may improve identification of interactions with previously unknown or related proteins.

AAC was included in this study however there are limitations with this type of representation. Using protein language models could be an area of development.

#### 2.9. Applicability Domain

The applicability domain (AD) defines the region of chemical and biological space in which the model can be expected to give reasonable results.

If the compounds and targets in the model are well represented in the training set then the model will be more reliable. Predictions made on compounds outside of the training chemistry may not be reliable. *(Tropsha, 2010)*

**There are many ways to identify the applicability domain:**
Molecule similarity (fingerprints)
Distance to training compounds
Outliers
Probability of prediction
Distribution shifts

Applicability domain is important in drug-target interaction prediction as novel chemical entities in external NCE libraries may not be well represented in historical bioactivity datasets.

A high predicted interaction score should not be assumed to have experimental confirmation. Predictions should be considered based on chemical similarity, coverage of the feature space and uncertainty.

The decrease in performance seen with the external NCE dataset shows the importance of including applicability domain in future iterations of this model.

#### 2.10 Gap in Research

Many DTI benchmarks in the literature only report performance on a limited number of targets using random splits. Often the label definitions, property bias and limitations of protein representations are not investigated. We provide a large random split evaluation (84,394 test records), a stringent external evaluation on NCEs removed from BindingDB, compare 5 ligand fingerprints under identical conditions, include physicochemical and ADMET context and openly report the generalisation gap and likely causes.

### 3. Research Objectives

#### 3. Research Objectives

The aim of this project is to develop and critically validate a machine-learning model that predicts potent drug–target binding from BindingDB Ki data. The specific objectives are:

1. To build a clean, reproducible Ki-based dataset from the BindingDB master file.
2. To study how the activity threshold changes class balance and to select a defensible cut-off.
3. To compare ligand representations (MACCS, ECFP4, ECFP6, Atom Pair and Topological Torsion) in one identical learning framework and identify the best performers.
4. To test whether physicochemical descriptors and nine predicted ADMET properties add useful context.
5. To measure generalisation on NCEs never seen during training and to explain any performance drop.
6. To describe how the model can support real drug discovery projects.

### 4. Methods

### 4. Methods

#### 4.1 Overview of the Workflow

The workflow (Figure 2) runs from the raw BindingDB release through filtering, labelling and feature construction to model training, threshold tuning and evaluation on internal and external data.

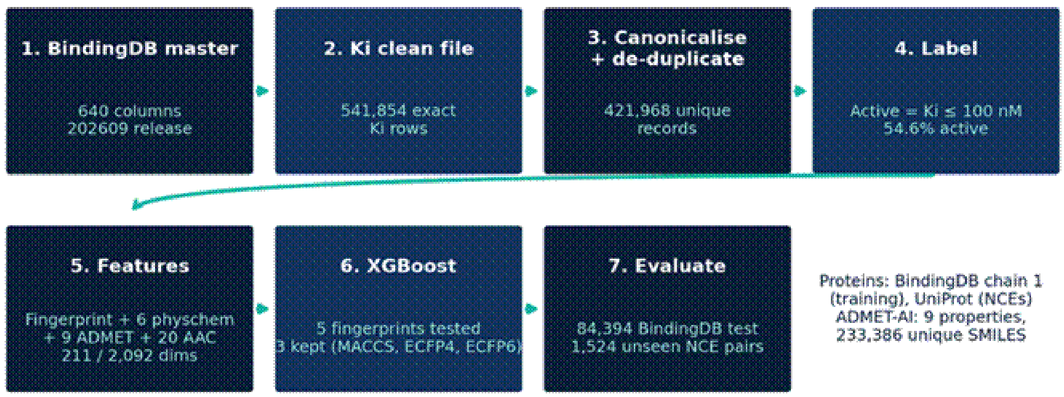

**Figure 2.** End-to-end workflow of the project with the record counts at each stage.

#### 4.2 BindingDB Master File and Filtering

We downloaded BindingDB_All_202609 (TSV), which has 640 columns, including per-chain fields for up to 50 protein chains. To keep memory manageable the file was streamed in 100,000-row chunks and only four fields were retained (Table 3).

**Table 3.** Fields retained from the BindingDB master file

| Retained field | Source column | Purpose |
|---|---|---|
| SMILES | Ligand SMILES | Ligand structure |
| Target_ID | Target Name | Target identity |
| Protein_Sequence | BindingDB Target Chain Sequence 1 | Protein features |
| Ki_nM | Ki (nM) | Source of the activity label |

Cleaning proceeded as follows. Rows missing SMILES, target, sequence or Ki were dropped. Only exact numeric Ki values were kept; qualified values such as ">10000" were converted to missing and removed, giving **541,854 rows** in BindingDB_Ki_clean.csv. SMILES were canonicalised with RDKit and molecules that failed parsing (for example pentavalent nitrogen) were removed. Records were de-duplicated on canonical SMILES plus protein sequence, giving **421,968 unique records**. An earlier prototype, de-duplicated on SMILES plus target name, contained 406,789 rows and was used for the first baselines.

#### 4.3 Protein Sequence Retrieval

Training proteins were taken directly from the BindingDB chain-sequence column. For the NCE set, target accessions were mapped to sequences through the UniProt REST FASTA service. In the first pilot 228 of 231 accessions were retrieved (P94451, Q7ZJM1 and Q9YQ12 failed). The final run retained 1,524 compound–target pairs with sequences.

#### 4.4 Ki Thresholding

Defining an active compound is the most consequential modelling decision. We evaluated three strict cut-offs on the 406,789-row prototype set and then adopted a more permissive one for the final models (Table 4, Figure 3).

**Table 4.** Effect of the Ki cut-off on class balance

| Ki cut-off | Active | Inactive | Active share |
|---|---|---|---|
| < 10 nM | 111,282 | 295,507 | 27.4% |
| < 30 nM | 156,806 | 249,983 | 38.5% |
| < 50 nM | 180,142 | 226,647 | 44.3% |
| ≤ 100 nM (final) | 295,855 | 245,999 | 54.6% (541,854 rows) |

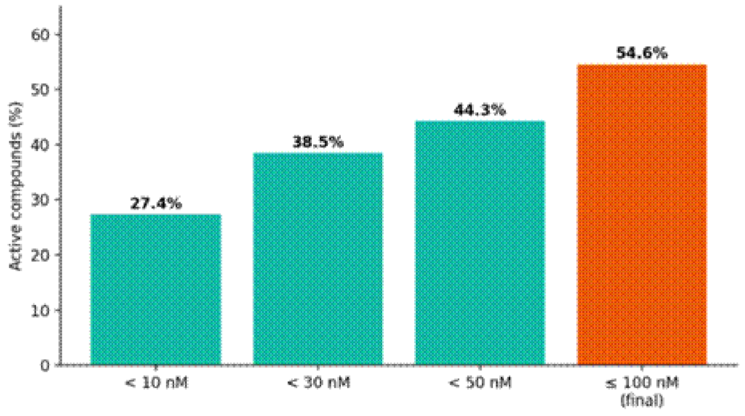

**Figure 3.** Share of active compounds under each Ki cut-off.

We chose **active = Ki ≤ 100 nM** for three reasons. First, 100 nM is a common potency criterion for lead-like, high-affinity binders. Second, a 10 nM cut-off leaves only 27.4% positives (about 1 : 2.7), giving the model fewer positive examples to learn from. Third, 100 nM yields a nearly balanced problem (54.6% active overall and 53.1% in the final test set), which makes threshold-dependent metrics such as F1 and MCC stable and meaningful.

#### 4.5 Feature Engineering

Each record is described by four feature blocks (Table 5): a ligand fingerprint, six physicochemical descriptors, nine ADMET properties and the amino-acid composition (AAC) of the protein. Fingerprint blocks were stored as sparse matrices.

**Table 5.** Feature blocks used for model training

| Block | Dimension | Details |
|---|---|---|
| Ligand fingerprint | 167 / 2,048 | MACCS keys; ECFP4 (Morgan radius 2); ECFP6 (Morgan radius 3) |
| Physicochemical | 6 | MW, Crippen logP, TPSA, H-bond donors, H-bond acceptors, QED |
| ADMET properties | 9 (+9 missing flags) | Nine ADMET-AI predicted properties, each with a missing-value indicator |
| Protein | 20 | Amino-acid composition |
| Total | 211 (MACCS); 2,092 (ECFP4/6) | Includes the nine missing-value flags |

**ADMET-AI integration.** ADMET-AI was installed from its GitHub repository. Throughput testing showed that batching matters: 32 molecules took 6.1 s in one batch, whereas predicting 128 molecules one at a time took 114 s (0.89 s per molecule). The final run predicted all 233,386 unique SMILES in batches. We selected nine properties that are relevant to exposure and safety: Caco-2 permeability, human intestinal absorption (HIA), plasma-protein binding (PPBR), CYP3A4 inhibition, CYP2D6 inhibition, hepatocyte clearance, half-life, hERG and drug-induced liver injury (DILI). Missing values were imputed with training-set medians only, which avoids leakage, and a binary missingness flag was added for each property.

#### 4.6 Baseline Models

A class-balanced random forest (300 trees) on MACCS plus AAC reached ROC-AUC 0.910 and AUPRC 0.798 on the 406,789-row prototype (Ki < 10 nM). XGBoost on ECFP4 (1,024 bits) plus AAC was then trained on the 541,854-row set. The two systems differ in both algorithm and fingerprint, so Table 6 compares pipelines and not an isolated algorithm.

**Table 6.** Random forest versus XGBoost baselines

| Metric | Random forest (MACCS + AAC) | XGBoost (ECFP4 + AAC) | Change |
|---|---|---|---|
| Accuracy | 0.8550 | 0.8414 | −0.0136 |
| Precision | 0.7440 | 0.6836 | −0.0604 |
| Recall | 0.7164 | 0.8309 | +0.1145 |
| F1 | 0.7299 | 0.7501 | +0.0202 |
| ROC-AUC | 0.9084 | 0.9195 | +0.0111 |
| AUPRC | 0.8071 | 0.8384 | +0.0313 |

XGBoost gave better ranking quality (AUC, AUPRC) and much higher recall, which is what a screening tool requires, at the cost of precision. It was carried forward.

#### 4.7 Final Model Configuration

- **Fingerprint screening:** MACCS, ECFP4, ECFP6, Atom Pair and Topological Torsion were tested under the same framework to find the best output; the three best were retained for the full study.
- **Split:** stratified 80% training-validation and 20% test (random state 42); 10% of the training portion was held out for early stopping, giving about 303.8k training, 33.8k validation and 84.4k test records (test: 39,570 inactive and 44,824 active).
- **Model:** XGBoost, histogram tree method, 1,500 trees, learning rate 0.03, maximum depth 8, row and column subsampling 0.8, scale_pos_weight from the training class ratio and early stopping after 100 rounds on validation AUC.
- **Decision threshold:** swept from 0.30 to 0.68 on the validation set only (selected on validation accuracy): 0.48 for MACCS and ECFP4 and 0.46 for ECFP6. The test set was never used for tuning.

#### 4.8 External NCE Dataset

The external set was built from a curated NCE compound table with ChEMBL identifiers, target accessions and pChEMBL values. After SMILES canonicalisation 7,319 molecules were valid. Every molecule whose canonical SMILES appears in BindingDB was removed (1,057 exact overlaps), leaving 6,262 compounds not found in BindingDB. Compounds with a pChEMBL value were mapped to a target accession and UniProt sequence. A pair was labelled active if pChEMBL $\geq$ 6.0 (potency $\leq$ 1 μM). The final evaluation set contains **1,524 compound–target pairs**, featurised with the same pipeline and evaluated at each model's validation-selected threshold.

#### 4.9 Evaluation Metrics

We report ROC-AUC, PR-AUC, accuracy, F1 score and the Matthews correlation coefficient (MCC), together with confusion matrices, sensitivity and specificity. PR-AUC and MCC are treated as primary metrics because they are robust to class imbalance and use more of the confusion matrix than accuracy.

#### 4.10 Development Lessons

Our first ADMET pilot used a small subset (2,841 rows) and reported accuracy near 0.99 but MCC of 0.00 to 0.51. Diagnosis showed that the label filter had retained only rows with raw Ki of 0 or 1 nM, so 99% of the data was positive and the model could score well by predicting active for everything. We corrected this by binarising Ki properly on the full dataset and by treating MCC and PR-AUC as primary metrics. The episode shows why a high accuracy alone should never be trusted.

### 5. Results and Discussion

### 5. Results and Discussion

#### 5.1 Results on BindingDB Test Set

The results from using the 3 selected fingerprints on the 84,394 datapoints in the test set can be seen in Table 7.

**Table 7.** Results on BindingDB Holdout Set.

| Fingerprint | Threshold | ROC-AUC | PR-AUC | Accuracy | F1 | MCC |
|---|---|---|---|---|---|---|
| MACCS | 0.48 | 0.9075 | 0.9158 | 0.8270 | 0.8391 | 0.6524 |
| ECFP4 | 0.48 | 0.9093 | 0.9189 | 0.8280 | 0.8386 | 0.6545 |
| ECFP6 | 0.46 | 0.9125 | 0.9222 | 0.8315 | 0.8440 | 0.6613 |

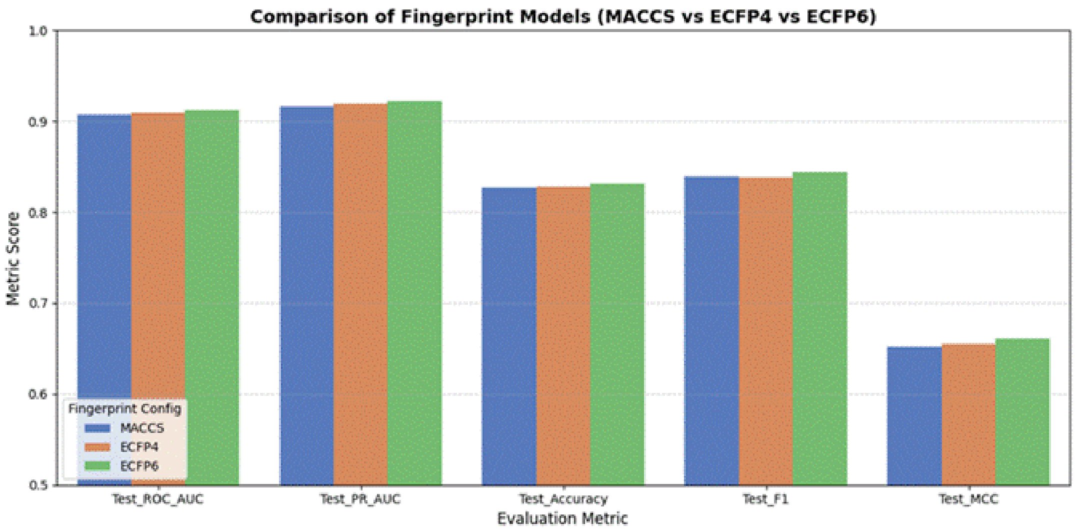

**Figure 4.** A comparison of MACCS, ECFP4 and ECFP6 using the 5 metrics

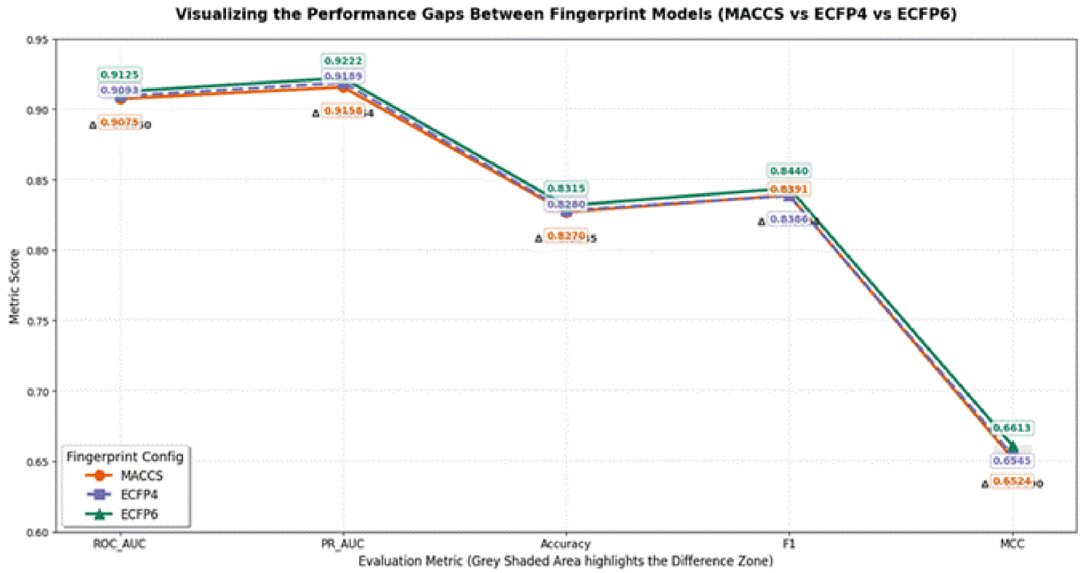

**Figure 5.** Difference in performance between the three fingerprint models. Exact scores have been included.

ECFP6 had the highest scores in all key metrics. However the differences between performances were small, with only 0.0050 difference in ROC-AUC, 0.0064 difference in PR-AUC, 0.0045 difference in ACCURACY and 0.0090 difference in MCC. Ligand features do not appear to be as important as having large amounts of data, along with protein/ADMET features. MACCS (167 bits) performs very similarly to the 2048 bit circular fingerprints meaning it would be a good choice if memory/speed is an issue. No early stopping was used since the AUC was still increasing. The validation AUC for MACCS, ECFP4 and ECFP6 at the 1500 th tree was 0.9056, 0.9075 and 0.9115 respectively.

#### 5.2 Confusion Matrices

**Table 8.** Confusion Matrix results on the BindingDB test set (84,394 datapoints)

| Model | TN | FP | FN | TP | Sensitivity | Specificity | Precision |
|---|---|---|---|---|---|---|---|
| MACCS | 31741 | 7829 | 6768 | 38056 | 0.849 | 0.802 | 0.829 |
| ECFP4 | 32164 | 7406 | 7110 | 37714 | 0.841 | 0.813 | 0.836 |
| ECFP6 | 31693 | 7877 | 6344 | 38480 | 0.859 | 0.801 | 0.830 |

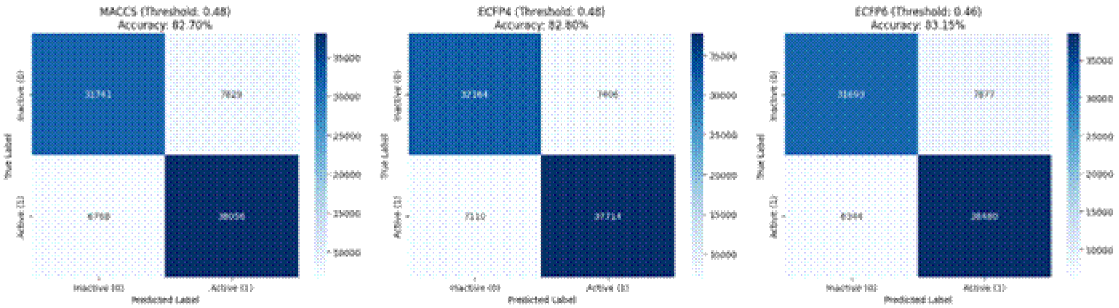

**Figure 6:** Confusion Matrices for MACCS, ECFP4 and ECFP6 models using the threshold selected in Validation.

ECFP6 has captured the most actives (38,480 true positives) and missed the least amount of actives(6,344). The models are not just predicting one class since all have a sensitivity between 0.84-0.86 and specificities between 0.80-0.81.

#### 5.3 Comparison of Fingerprints

The results in table 9 show that Atom Pair and Topological Torsion both performed worse than MACCS in all metrics. They also are not identical to MACCS. They used vectors of 2,048 bits compared to MACCS which used 167 bits. They therefore provided no benefit to using at a higher cost so MACCS (substructure keys) and ECFP4/ECFP6 (circular fingerprints) were used.

**Table 9.** Comparison of Models for Test fingerprints

| Rank | Fingerprint | Bits | ROC-AUC | PR-AUC | Accuracy | F1 | MCC | Decision |
|---|---|---|---|---|---|---|---|---|
| 1 | ECFP6 | 2048 | 0.9125 | 0.9222 | 0.8315 | 0.8440 | 0.6613 | Selected (Best) |
| 2 | ECFP4 | 2048 | 0.9093 | 0.9189 | 0.8280 | 0.8386 | 0.6545 | Selected |
| 3 | MACCS | 167 | 0.9075 | 0.9158 | 0.8270 | 0.8391 | 0.6524 | Selected |
| 4 | Atom Pair | 2048 | $< 0.9075$ | $< 0.9158$ | $< 0.8270$ | $< 0.8391$ | $< 0.6524$ | Performed worse than MACCS |
| 4 | Topological Torsion | 2048 | $< 0.9075$ | $< 0.9158$ | $< 0.8270$ | $< 0.8391$ | $< 0.6524$ | Performed worse than MACCS |

#### 5.4 External Validation on Unseen NCEs

**Table 10.** Results on unseen NCEs (1524 pairs)

| Fingerprint | Threshold | ROC-AUC | PR-AUC | Accuracy | F1 | MCC |
|---|---|---|---|---|---|---|
| MACCS | 0.48 | 0.7895 | 0.8232 | 0.7106 | 0.7249 | 0.4297 |
| ECFP4 | 0.48 | 0.7884 | 0.8302 | 0.7041 | 0.7187 | 0.4164 |
| ECFP6 | 0.46 | 0.8040 | 0.8464 | 0.7316 | 0.7464 | 0.4701 |

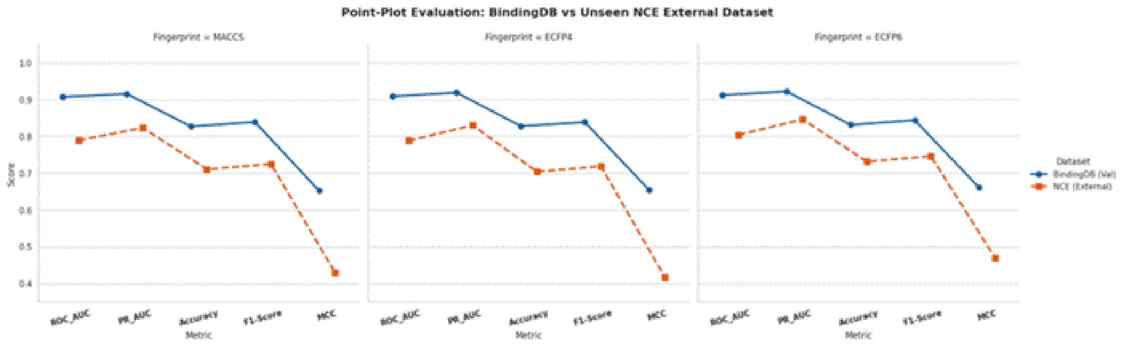

**Figure 7.** Point plot comparing performance of each fingerprint on BindingDB Validation and Unseen NCEs.

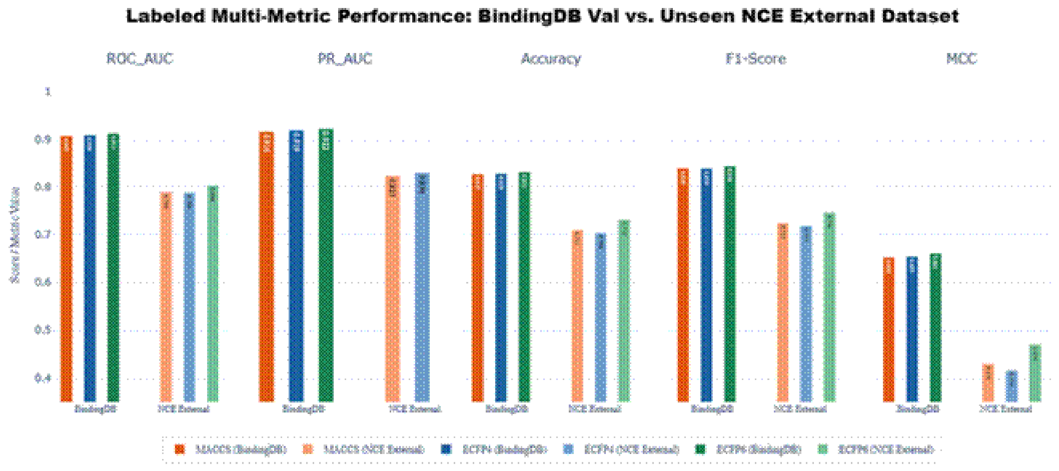

**Figure 8**: Multi-metric comparison of model predictions (BindingDB Test vs Unseen NCEs). Includes labels.

ECFP6 still performs well on truly unseen compounds, achieving a ROC-AUC score of 0.8040, PR-AUC score of 0.8464 and MCC score of 0.4701. Earlier work with ECFP4 model found that with a smaller set of 228 NCE pairs, the ROC-AUC score was 0.735, PR-AUC score 0.681 and accuracy score 0.71. Whilst inactives were well recovered (recall score 0.87), actives were not predicted well (recall score 0.47). The larger size of the final set will give more reliable results.

#### 5.5 Generalisation Gap

**Table 11.** Differences in performance between BindingDB test set and NCE set

| Metric | MACCS | ECFP4 | ECFP6 |
|---|---|---|---|
| ROC-AUC | 0.1180 | 0.1209 | 0.1085 |
| PR-AUC | 0.0926 | 0.0887 | 0.0758 |
| Accuracy | 0.1<br>164 | 0.1239 | 0.0999 |
| F1 | 0.1142 | 0.1199 | 0.0976 |
| MCC | 0.2227 | 0.2381 | 0.1912 |

ECFP6 had the smallest drop in all 5 metrics. Having a radius of 3 bonds provides more contextual information and seems to provide a more generalizable model than using 167 predetermined keys or a radius of 2 Hashes.

#### 5.6 Factors that contribute to drop in model performance

The reasons for the drop in external NCE performance can be attributed to:

- External NCEs were not present within the training set from BindingDB. Random splitting may have resulted in similar analogues being present within the training and test set, therefore increasing the internal performance.

- There was a large discrepancy in activity thresholds between the training set (Ki ≤ 100nM) and testing set (pChEMBL ≥ 6, ≤ 1μM). This is a 10 fold difference and the variety of assays could have impacted classification performance.

- NCEs did not have any available ADMET data. Therefore these were replaced with median values from training set. These were also flagged as missing data, however this was not often seen during training.

- AAC has only 20 features so sequence order and binding site information cannot be captured. This limits the ability to distinguish between similar proteins.

Class imbalance and Threshold Differences could have resulted in reduced external classification.

Overall, this shows that there are differences between the internal and external datasets which caused the drop in performance, however it does not necessarily indicate that the model has failed fundamentally. The model still performed well in terms of ROC-AUC (0.8040) and PR-AUC (0.8464) when tested on unseen NCEs showing it can prioritise potentially active compounds.

### 6. Conclusion and Future Scope

### 6. Conclusion and Future Scope

#### 6.1 Summary of Findings

The XGBoost-based DTI prediction framework developed has shown promising results using a combination of molecular fingerprints, physiochemical properties, ADMET properties and protein amino acid composition.

Findings include:

Data Preparation – 421,968 unique interactions were obtained from the processed BindingDB dataset. Compounds with a Ki value of under or equal to 100nM were classed as active.

Model Performance – Using a combination of features, the XGBoost model achieved strong results with a ROC-AUC of 0.9125 and MCC of 0.66 on 84,394 test instances.

Molecular Fingerprints – When evaluating the five molecular fingerprints, the ECFP6 fingerprints achieved the best results. The MACCS fingerprints had similar results with only a difference in ROC-AUC of ~0.005. The Atom Pair and Topological Torsion fingerprints had poorer results.

Model Performance (NCE) – On evaluation of the model with 1524 NCE interactions that had not been seen the model achieved a ROC-AUC of 0.8040, PR-AUC of 0.8464 and MCC of 0.4701. This shows there was a drop in performance when evaluating unknown compounds.

#### 6.2 Benefits of the Proposed Framework

The framework has strong ROC-AUC results both internally and externally which shows it has potential to identify promising drug-target interactions.

This framework has fast screening capabilities as all the molecular and protein features can be calculated computationally. The ADMET-AI feature took ~0.2 seconds per molecule.

Developability – The model includes nine ADMET properties to give further insight into how well they will be absorbed, metabolised and any potential toxicities.

Large-scale screening – The framework allows for large scale screening with flexibility on the different molecular fingerprints.

This framework has potential to improve early stage drug discovery due to the computational prioritisation of compounds.

By combining the two frameworks together, the lower priority candidates can be filtered out so resources can be focused on the compounds that have favourable predictions for interactions and developability.

This may improve the efficiency of hit identification and hit-to-lead optimisation. However, there is no measurement of how much faster the development would be or how much less it would cost to run experiments.

**Practical example.** Consider a team with a series of new analogues against one protein target. (1) For each analogue the SMILES and the target UniProt sequence are entered. (2) The pipeline computes the ECFP6 fingerprint, six physicochemical descriptors, nine ADMET properties and the protein composition. (3) The XGBoost model returns a probability of Ki ≤ 100 nM. (4) A probability of at least 0.46 flags the compound as likely potent. (5) Flagged compounds with acceptable ADMET predictions go to the assay, and the rest are deprioritised before synthesis or purchase. On unseen NCEs, an ROC-AUC of 0.8040 means an active compound is ranked above an inactive one about 80% of the time.

#### 6.3 Our View

Our strongest contributions are a reproducible, memory-aware pipeline for a 500k-row dataset, a controlled fingerprint comparison, validation-only threshold selection, an external test with explicit overlap removal and transparent reporting of the generalisation gap and of a bug found and fixed. We do not claim that 83% internal accuracy predicts prospective performance, that ADMET features were proven useful on the NCEs, or that the fingerprint ranking is statistically settled, since differences are small and based on a single random seed.

#### 6.4 Future Scope

**Table 12.** Proposed next steps

| No. | Next step | Expected benefit |
|---|---|---|
| 1 | Run ADMET-AI on all NCE compounds and re-evaluate | Removes the feature mismatch noted in Section 5.6 |
| 2 | Align NCE labels with the training definition (Ki, ≤ 100 nM) or retrain at pChEMBL ≥ 6 | Aligns label semantics |
| 3 | Scaffold-based, cluster-based and target-held-out splits | More honest internal estimates |
| 4 | Replace AAC with learned protein embeddings such as ESM-2 | Target-specific signal |
| 5 | Ablations: fingerprint only, then physicochemical, ADMET and protein blocks added | Shows each block's contribution |
| 6 | Multiple seeds, confidence intervals and significance tests | Settles the fingerprint ranking |
| 7 | More trees or Bayesian hyperparameter search; ECFP6 + MACCS ensembles | Uses unconverged capacity |

| | | |
|---|---|---|
| 8 | Probability calibration, applicability-domain scoring and SHAP interpretation | Trustworthy, explainable predictions |
| 9 | pKi regression and multi-task learning with Kd and IC50 | More data and potency estimates |
| 10 | Prospective validation: rank top NCEs, then docking or wet-lab assays | Real-world proof |

---

**Data and software links** (not in the report's reference list, but worth citing in Methods):

- BindingDB: https://www.bindingdb.org
- ChEMBL: https://www.ebi.ac.uk/chembl/
- UniProt: https://www.uniprot.org
- PubChem: https://pubchem.ncbi.nlm.nih.gov
- ADMET-AI code: https://github.com/swansonk14/admet_ai
- XGBoost documentation: https://xgboost.readthedocs.io
